# Characterization of ultrapotent chemogenetic ligands for research applications in non-human primates

**DOI:** 10.1101/2022.01.06.475241

**Authors:** Jessica Raper, Mark A. G. Eldridge, Scott. M. Sternson, Jalene Y. Shim, Grace P. Fomani, Barry J. Richmond, Thomas Wichmann, Adriana Galvan

## Abstract

Chemogenetics is a technique for obtaining selective pharmacological control over a cell population by expressing an engineered receptor that is selectively activated by an exogenously administered ligand. A promising approach for neuronal modulation involves the use of “Pharmacologically Selective Actuator Modules” (PSAMs); these chemogenetic receptors are selectively activated by ultrapotent “Pharmacologically Selective Effector Molecules” (uPSEMs). To extend the use of PSAM/PSEMs to studies in nonhuman primates it is necessary to thoroughly characterize the efficacy and safety of these tools. We describe the time course and brain penetrance in rhesus monkeys of two compounds with promising binding specificity and efficacy profiles in *in vitro* studies, uPSEM792 and uPSEM817, after systemic administration. Rhesus macaques received subcutaneous (s.c.) or intravenous (i.v.) administration of uPSEM817(0.064 mg/kg) or uPSEM792 (0.87 mg/kg) and plasma and CSF samples were collected over the course of 48 hours. Both compounds exhibited good brain penetrance, relatively slow washout and negligible conversion to potential metabolites - varenicline or hydroxyvarenicline. In addition, we found that neither of these uPSEMs significantly altered heart rate or sleep. Our results indicate that both compounds are suitable candidates for neuroscience studies using PSAMs in nonhuman primates.

## BACKGROUND

The most widely used chemogenetic system is currently the hM_3_Dq/hM_4_Di DREADDs, which are based on G-protein coupled receptors (Armbruster et al., 2007). However, DREADDs have several drawbacks. Since they work via second messangers, they have multiple mechanisms of action to facilitate or attenuate neuronal firing (Atasoy and Sternson, 2018). Another drawback is that, despite there being several ligands to activate DREADDs, each has at least some degree of off-target binding(Ilg et al., 2018; MacLaren et al., 2016; Manvich et al., 2018; Roseboom et al., 2021; Upright and Baxter, 2020). Another chemogenetic approach is based on engineered Ligand-Gated Ion Channels (LGICs).These ion channels directly influence the electrical properties of cell membranes using engineered “Pharmacologically Selective Actuator Modules” (PSAMs). These are channels that are chimeric LGICs comprising a mutated α7 nicotinic acetylcholine receptor (α7-nAChR) ligand binding domain, combined with the ion pore domain of either the excitatory cation-selective 5-HT3a receptor, or the inhibitory chloride-selective glycine receptor (GlyR).(Sternson and Roth, 2014) PSAMs are selectively activated by ultrapotent “Pharmacologically Selective Effector Molecules” (uPSEMs). We have previously shown that systemic administration of varenicline, an FDA-approved agonist for one class of PSAMs, reduces neuronal activity in neurons expressing PSAM-GlyRs in rhesus monkeys (Magnus et al., 2019).

Here, in rhesus monkeys we measured the time course and brain penetrance of two new ultrapotent PSEM compounds, uPSEM792 and uPSEM817, that show promising binding specificity and efficacy profiles in *in vitro* studies (Magnus et al., 2019). We also analyzed metabolite concentrations and monitored the occurrence of side effects associated with the use of these compounds by tracking physical activity, sleep, and heart rate. Overall the PSEMs were well tolerated.

## METHODS

Six rhesus monkeys were used in this study; two from the National Institute of Mental Health (one 7-year-old female weighing 10.4 kg, and one 13-year-old male weighing 11.4 kg at the beginning of studies) and four from the Yerkes National Primate Research Center (males, 4-5 years old and weighing 6.5-7.2 kg).

All monkeys were pair-housed indoors on a 12 hour light/dark cycle, had free access to food and water, and received vegetables and fruit daily. All procedures conformed to the *Institute for Laboratory Animal Research* Guide and were performed under an Animal Study Proposal approved by the Animal Care and Use Committee of the National Institute of Mental Health or the Institutional Animal Care and Use Committee of Emory University, and were in accordance with the United States Department of Agriculture (USDA) Animal Welfare Act.

### Drugs and formulation

The uPSEM doses were selected based on the lowest effective doses in mice (Magnus et al., 2019) and excluded the mass of the counterion (see below). uPSEM817 tartrate was dissolved in sterile saline, acidified to pH 4-5 with HCl. The acidification step was not followed when the uPSEM817 solution was prepared at Yerkes. Sterile saline was added to bring the concentration of the solution to 1 mg/mL, and the pH then titrated back to neutral (pH 6 – 7) with NaOH. The solution was passed through a 0.22-micron sterile filter prior to administration at a dose of 0.064 mg/kg (0.1 mg/kg before correction for the tartrate counterion). uPSEM792 hydrochloride was formulated and filtered per the protocol described above, and was administered at a dose of 0.87 mg/kg (1.0 mg/kg before correction for the chloride counterion). The compounds used were obtained from Dr. Sternson’s lab (Janelia), Tocris (MN) or HelloBio (NJ).

### uPSEM Intravenous Administration

Intravenous administration challenges with uPSEM817 and uPSEM792 were done in two monkeys.

#### Surgeries

The macaques were implanted with indwelling subcutaneous access ports (Access Technologies, Illinois) for the collection of blood and cerebrospinal fluid (CSF). The blood collection ports were connected to a catheter implanted into the femoral artery using a procedure similar to that previously described for femoral vein catheterization (Graham et al., 2008). The ports were flushed with saline and locked with taurolidine citrate solution (TCS, Access Technologies, Illinois). The CSF collection ports were implanted using a variation on a previously described technique (Felice et al., 2011), instead of a hemilaminectomy for access to the thecal sac, a laminotomy was performed. The CSF access port was flushed with saline and locked with dilute heparinized saline (10 USP/mL).

#### Intravenous uPSEM injections and sample collection

Baseline blood and CSF samples were obtained from the chronically-implanted subcutaneous access ports. To clear the ‘dead space’ in the blood access port, 0.5 mL blood was extracted and discarded. A sample of 2 mL of blood was then obtained and transferred to a 2 mL tube containing EDTA (3.5 mg) on ice. To clear the ‘dead space’ in the CSF access port, 0.25 mL CSF was withdrawn and discarded. A 0.5 mL sample of CSF was collected and immediately transferred to dry ice. Either uPSEM817 (0.064 mg/kg) or uPSEM792 (0.87 mg/kg) was injected 30 min after baseline sample collection via a temporary catheter in the saphenous vein. Further blood and CSF samples were collected, per the procedure used for baseline collections, at the following time points post i.v. drug injection: 10, 20, 40, 60, 90, 120, 180, 360, 720 min, 24 hr, and 48 hr. All blood samples were centrifuged at 2500 rcf for 10 min within 60 min from time of collection, and the plasma was transferred to Eppendorf tubes for storage at −80°C. CSF samples were transferred directly to the −80°C freezer without thawing.

### Subcutaneous uPSEM administration

Three unoperated adult rhesus were used in uPSEM817 subcutaneous administration challenges. For uPSEM817, the three monkeys received a subcutaneous injection of uPSEM817 (0.064 mg/kg) and were then sedated with tiletamine HCL and zolazepam (Telazol, 3-5mg/kg, im). CSF samples were obtained in a serial fashion (as explained below) from the cisterna magna, using a sterile 23-ga bevel-tipped needle by pressure difference and collected by gravity (Raper et al., 2017). CSF samples were collected in pre-chilled sterile Eppendorf tubes, immediately frozen on dry ice, and stored at −80 °C until the time of the assay.

Serial CSF taps were performed in sessions separated by 2 weeks. To reduce the risks associated with repeated spinal taps and given the time restrictions imposed by the anesthetized state, in each session, only 2-3 serial CSF taps were conducted per animal, so that not all time points were collected for all monkeys. Thus, after subcutaneous injection of uPSEM817 0.064 mg/kg, three individual CSF samples were available for analyses at 15, 30, 60, 90, 120, 1440, 2880 min post-injection, and two individual CSF samples at 5, 45, and 240 min post injection.

Blood samples were collected from the femoral vein immediately following each CSF tap. All blood samples were collected in pre-chilled 2 mL tubes containing EDTA (3.5 mg) and immediately placed on ice. Samples were centrifuged at 2500 rcf for 15 min in a refrigerated centrifuge (at 4 °C). Plasma was pipetted off and stored at −80 °C until assayed.

The same two adult rhesus macaques used in the i.v. administration were also used for the subcutaneous administration challenge of 0.87 mg/kg dose of uPSEM792. All procedures were the same as described earlier. The post s.c. drug injection samples were collected at the same time points as uPSEM817 post s.c. drug collections.

### Microdialysis

In one monkey, microdialysis experiments were conducted to measure the concentrations of uPSEM792 or uPSEM817 in the putamen before and after systemic injections of these compounds. Under isoflurane anesthesia and sterile conditions, the animal received a chronic recording chamber directed to the putamen, following previously described procedures (Bogenpohl et al., 2013; Nanda et al., 2009). Prior to the microdialysis experiments, the putamen was identified with electrophysiological mapping, according to its stereotaxic location, depth in the dorso-ventral plane, its relationship with other structures as well its characteristic neuronal firing activity (Bogenpohl et al., 2013; DeLong, 1973; Nanda et al., 2009).

Thereafter, microdialysis probes (modified CMA-11, CMA Microdialysis, Kista, Sweden, 3 mm cuprophane membrane, molecular weight cut-off, 6 kD) were inserted into the putamen through a 22-gauge guide cannula. Two microdialysis experiments were performed, one each for uPSEM792 and uPSEM817. The probes were perfused with artificial cerebrospinal fluid (aCSF, CMA) at 2 μl/min. After probe insertion, two baseline (before injection) samples were collected 60 min apart (120 μl). uPSEM817 (0.064 mg/kg) or uPSEM792 (1 mg/kg) was then injected s.c. Five minutes after the injection, we started the collection of the post-injection samples, and three samples were collected (one sample every 40 min, 80 ul/sample). Following the experiment, the probe was removed from the brain and placed into a vial with a standard uPSEM solution, and an additional sample was collected for 40 min. This sample was used to calculate the recovery fraction of each microdialysis probe. We report the the values corrected by this recovery fraction (11% and 7% for the uPSEM817 and uPSEM792 experiments respectively).

### Analysis of biological samples

All plasma, CSF, and microdialysis samples were assayed for the injected compound by Q2 solutions (Indianapolis, IN, see Supplemental Material for details). The plasma and CSF samples were also assayed for hydroxyvarenicline and varenicline. Both are potential metabolites of uPSEM817 and uPSEM792.(Obach et al., 2006) Hydroxyvarenicline can convert to varenicline, and varenicline can directly interact with a variety of endogenous receptors. Given the negligible levels of these compounds detected in CSF samples after the uPSEM817 or uSPEM792 injections (see below), the microdialysis samples were not assayed for metabolites.

### Heart Rate Monitoring

Three adult rhesus macaques (1 female, 2 males) were used to determine whether uPSEM817 and uPSEM792 had any effect on heart rate, as previously shown with drus that activate nAChRs (e.g. nicotine, varenicline.) (Donny et al., 2011; Jutkiewicz et al., 2013). Two monkeys were trained to voluntarily present a leg for heart rate monitoring via a wearable LED optical sensor (FitBit Inspire 2, San Francisco, CA or Polar OH1, Polar Electro Oy, Kempele, Finland) and one animal was monitored with an optical pulse-oximeter attached to its ear (SurgiVet® Advisor® Vital Signs Monitor by Smith Medical, Dublin, OH). For additional details see Supplemental Material”After recording baseline heart rate, the monkeys received a subcutaneous injection of either 0.064 mg/kg uPSEM817, 0.87 mg/kg uPSEM792, or vehicle (phosphate-buffered saline, Corning, Corning, New York). The heart rate was recorded immediately after injection, at 1 min post-injection, and at 5 min intervals for 30 min post-injection. Each animal’s heart rate was monitored in this manner 2-3 times for each compound (uPSEM817, uPSEM792, vehicle) with at least 1 week between uPSEM compound administrations.

### Sleep and Activity Monitoring

These experiments were only carried out with uPSEM817. To determine whether the agent affected sleep or activity levels, the agent was injected in two adult male monkeys wearing accelerometers (Actical, Respironics Inc., Murrysville, PA) that were attached to the collars. For five separate weeks, the monkeys’ activity levels were continuously measured from Friday to Monday morning. Accelerometers were placed on the monkeys’ collars on a Friday and the monkeys were given an s.c. injection of either 0.064 mg/kg uPSEM817 (3 separate sessions per monkey) or vehicle (PBS; 2 separate sessions per monkey). Accelerometry data were then analyzed with the standard Actiware algorithm (Respironics), (Paquet et al., 2007) which defines sleep on a weighted, sliding average with a threshold optimized for nighttime activity of nonhuman primates (the threshold for determining the onset of sleep was set to 14, instead of the standard setting of 20 (Cortes et al., 2016)) and sensitivity to detect night-time sleep disruption. Sleep efficiency was quantified as the total number of minutes slept each night as the percentage of the duration of the lights-off period (7 pm – 7 am), as described previously (Barrett et al., 2009; Cortes et al., 2016). The number of bouts of uninterrupted sleep and their average duration was also calculated. The latency to fall asleep was defined as time between lights off and the beginning of the first ≥3-min period of uninterrupted sleep. Daytime activity was calculated as the average activity/min starting when the animal was returned to their home cage on the day of injection until lights off (1 pm-7 pm). The same daytime activity period was compared for the day of injection, day 1 (24 hr post-injection) and day 2 (48 hr postinjection).

### Statistical Analyses

The area under the curve (AUC), maximal plasma concentration (Cmax), and time of maximal plasma concentration (Tmax), derived from plots of concentration over time, were calculated for uPSEM817, uPSEM792, varenicline, and hydroxyvarenicline, using GraphPad Prism 7.01 (GraphPad Software Inc. La Jolla, USA). The data are reported as means ± SEM. The molar mass used for calculations was 419.43 g/mol uPSEM817 tartrate salt, 277.75 g/mol for uPSEM792 hydrochloride, 211.26 g/mol for varenicline (Pubchem ID 5310966), and 227.26 g/mol for hydroxyvarenicline (M3b; Pubchem ID 29982201). The oil:water partition coefficient (cLogP) of uPSEM817 and uPSEM792 was calculated using Chem Draw Professional (PerkinElmer).

The change from baseline heart rate across the monitored post-injection time was calculated separately for the uPSEM817, uPSEM792, and vehicle experiments. Data were analyzed with a linear mixed model (LMM), with percent change from baseline heart rate as the outcome variable, drug injection (uPSEM817, uPSEM792, Vehicle) and time post-injection (0, 1, 5, 10, 15, 20, 25, 30min post) as fixed effects, and individual monkey as a random effect to account for repeated sessions for each monkey.

Sleep efficiency, latency to fall asleep, number of uninterrupted sleep bouts, average duration of sleep bouts, and daytime activity levels were analyzed using a LMM with drug injection (uPSEM817, Vehicle) and day post-injection (0=same day, 1, 2 days post) as fixed effects, and individual monkey as a random effect.

## RESULTS

### Pharmacokinetics of uPSEM817

The maximal plasma concentration was quickly reached after s.c. administration (Tmax = 15min). Note that the Tmax for the i.v. administrations technically occured at the time of administration. The Cmax was 71% higher after i.v. administration (11.78 vs 6.88 ng/mL for i.v. and s.c. administration, respectively; Table 1, Figure 1A). Similarly, the AUC value produced by i.v. administration (2.56 ng/(mL·h)) was 86% higher than that produced by s.c. administration (AUC, 1.38 ng/(mL·h); see Table 1). The uPSEM817 plasma concentration remained stable for approximately 3 hours post injection, and dropped below 0.5 ng/mL by 24 h.

**Table 1.** Pharmacokinetic Parameters of uPSEM817.

|  | Route | uPSEM817 |  | Hydroxyvarenicline |  |
| --- | --- | --- | --- | --- | --- |
|  |  | Plasma | CSF | Plasma | CSF |
| **AUC**<br>(ng/ml hr) | **s.c.** | 1.382 | 0.799 | 0.556 | 0 |
|  | **i.v.** | 2.564 | 1.154 | 1.161 | 0.0015 |
| **C<sub>max</sub>**<br>(ng/ml) | **s.c.** | 6.883 | 2.67 | 0.553 | NA |
|  | **i.v.** | 11.78 | 2.06 | 0.874 | 0.05 |
| **C<sub>max</sub> (nM)** | **s.c.** | 16.41 | 6.37 | 2.43 | NA |
|  | **i.v.** | 28.07 | 4.911 | 3.85 | 0.22 |
| **T<sub>max</sub>** | **s.c.** | 15 min | 30 min | 60 min | NA |
|  | **i.v.** | 0 min | 90 min | 180 min | 90 min |
| Mean plasma and cerebrospinal fluid (CSF) values for uPSEM817 (0.064 mg/kg) with administration route via subcutaneous (s.c.) or intravenous (i.v.) injection. AUC: area under the curve; C<sub>max</sub>: maximal concentration; T<sub>max</sub>: time at maximal concentration. Not Applicable (NA) indicates that hydroxyvarenicline was not detected in any of the samples. Parameters for varenicline are not listed, as this compound was not detected at any time point. |  |  |  |  |  |

**Figure 1.**
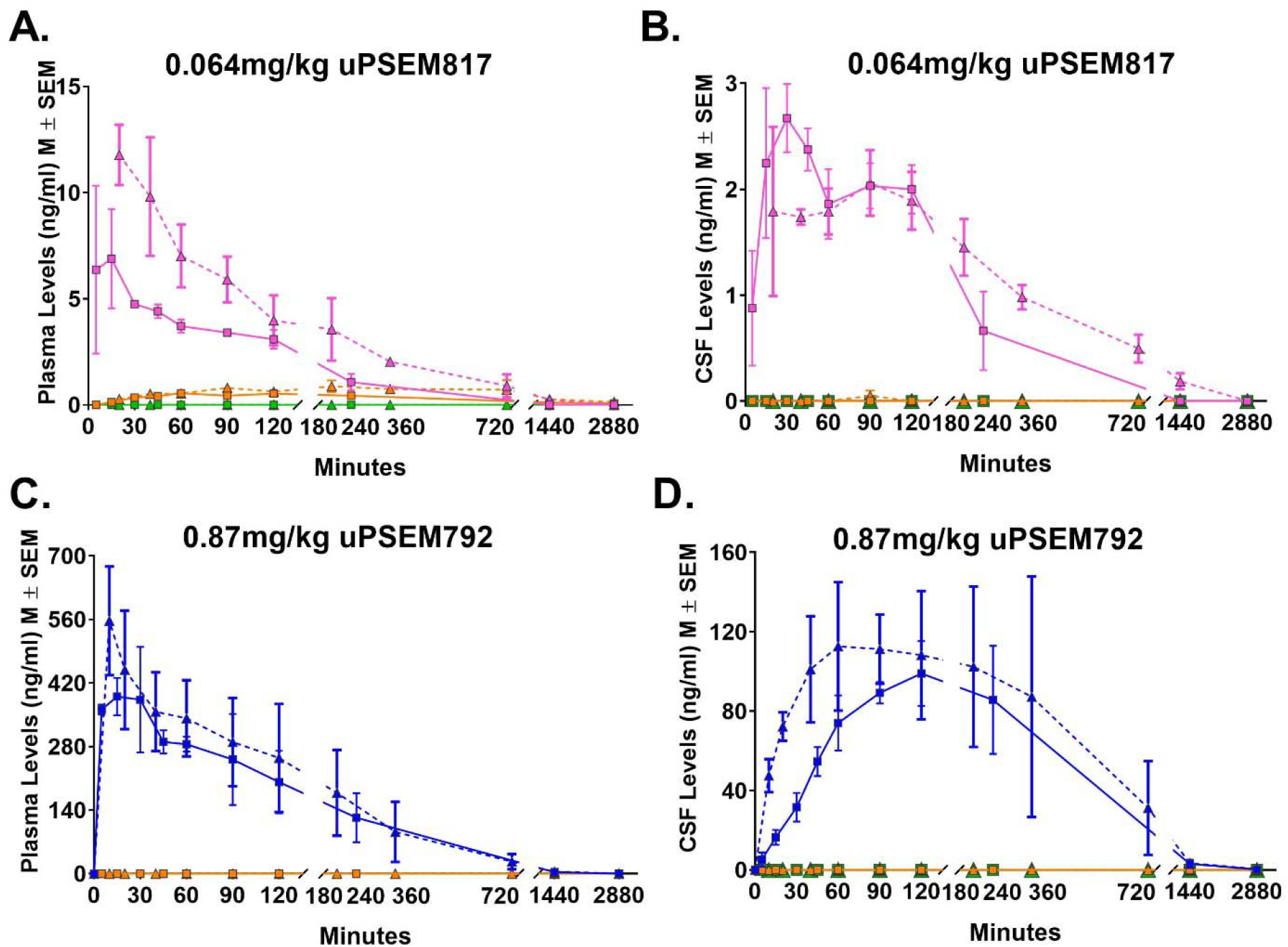
Time – concentration profiles of uPSEMs and metabolites. Levels of uPSEM817 (pink) and metabolites, hydroxyvarenicline (orange) or varenicline (green), in plasma (A) and CSF (B) after an intravenous (i.v.; triangle with dashed line) or subcutaneous (s.c.; square with solid line) administration of 0.064 mg/kg dose of uPSEM817. Panels C and D illustrate the levels of uPSEM792 (blue) and metabolites in plasma (C) and CSF (D) after i.v. or s.c. administration of 0.87 mg/kg dose of uPSEM792.

The maximal concentration in CSF (2.67 ng/mL, 6.37 nM) was reached within 30 min of the s.c. administration, whereas i.v. administration had a Tmax of 90 min (2.06 ng/ml, 4.91 nM; Table 1, Figure 1B). In the microdialysis experiment, we confirmed that uPSEM817 was not detectable in the brain before administration of uPSEM817. After s.c. injection, of uPSEM817, the concentrations of the compound in the brain microdialysates were 20.9, 20.1 and 8.2 ng/ml at 45, 85 and 125 min, respectively (corresponding to 49.8, 47.9 and 19.6 nM).

Following a single s.c. or i.v. administration of uPSEM817 (0.064 mg/kg), metabolic conversion of uPSEM817 to hydroxyvarenicline was detected in plasma with a very low Cmax of 0.55 ng/mL, 60 min after s.c. injections, and 0.87 ng/mL, 180 min after i.v. administration. Hydroxyvarenicline was only detected in a single CSF sample for one animal, 90 min post-injection for i.v. administration. Varenicline was not detected in either plasma or CSF, indicating that hydroxyvarenicline was not metabolized to varenicline.

### Pharmacokinetics of uPSEM792

We used a 14-fold higher dose of uPSEM792 than the dose used in *in vivo* studies in mice. The Tmax for plasma samples after s.c. administration (15 min) was similar to the Tmax for uPSEM817. The Cmax after s.c. administration (390.4 ng/mL) was 70% of the level measured after i.v. administration (557 ng/mL; Table 1, Figure 1C). However, the AUC values produced after s.c. administration were modestly higher than for i.v. administration (133.24 and 114.48 ng/(mL·h), respectively Table 2). The uPSEM792 plasma concentration slowly declined over approximately 6 h post injection and was still detectable at 48 h.

**Table 2.** Pharmacokinetic Parameters of uPSEM792.

|  | Route | uPSEM792 |  |
| --- | --- | --- | --- |
|  |  | Plasma | CSF |
| **AUC**<br>(ng/ml hr) | **s.c.** | 133.24 | 74.124 |
|  | **i.v.** | 114.48 | 71.342 |
| **C<sub>max</sub>**<br>(ng/ml) | **s.c.** | 390.4 | 98.95 |
|  | **i.v.** | 557.0 | 112.7 |
| **C<sub>max</sub> (nM)** | **s.c.** | 1405.6 | 356.26 |
|  | **i.v.** | 2005.4 | 405.76 |
| **T<sub>max</sub>** | **s.c.** | 15 min | 120 min |
|  | **i.v.** | 0 min | 60 min |
| Mean plasma and cerebrospinal fluid (CSF) values for uPSEM792 (0.87 mg/kg) with administration route via subcutaneous (s.c.) or intravenous (i.v.) injection. AUC: area under the curve; C<sub>max</sub>: maximal concentration; T<sub>max</sub>: time at maximal concentration. Hydroxvarenicline and varenicline were not detected at any time point. |  |  |  |

The maximal concentration in CSF (98.95 ng/mL, 356.26 nM) was reached within 120 min after s.c. administration, whereas i.v. administration resulted in a Tmax of 60 min (112.7 ng/ml, 405.76 nM; Table 2, Figure 1D). Collection of microdialysates in the putamen showed that uPSEM792 was not detectable in the brain prior to administration of the drug. After s.c. injection, the brain microdialysate concentrations of the compound were 41.9, 67.7 and 70.7 ng/ml at 45, 85 and 125 min, respectively, after injection (corresponding to 150, 243 and 254 nM).

Regardless of administration route (s.c. or i.v.), we did not observe any evidence for *in vivo* metabolism of uPSEM792 to hydroxyvarenicline or varenicline in either plasma or CSF (Figures 1C & D).

### Heart rate measurements

The analysis of the percent change from baseline heart rate revealed a significant main effect of time (F [7,102] = 17.96, p < 0.001), such that the heart rate increased immediately after the injections and slowly came back down toward baseline over the 30 min of monitoring. The change in heart rate was indistinguishable between uPSEM817 (0.064 mg/kg), uPSEM792 (0.87 mg/kg) and vehicle injections (Drug: F [2,102] = 0.097, p = 0.91; Figure 2A).

**Figure 2.**
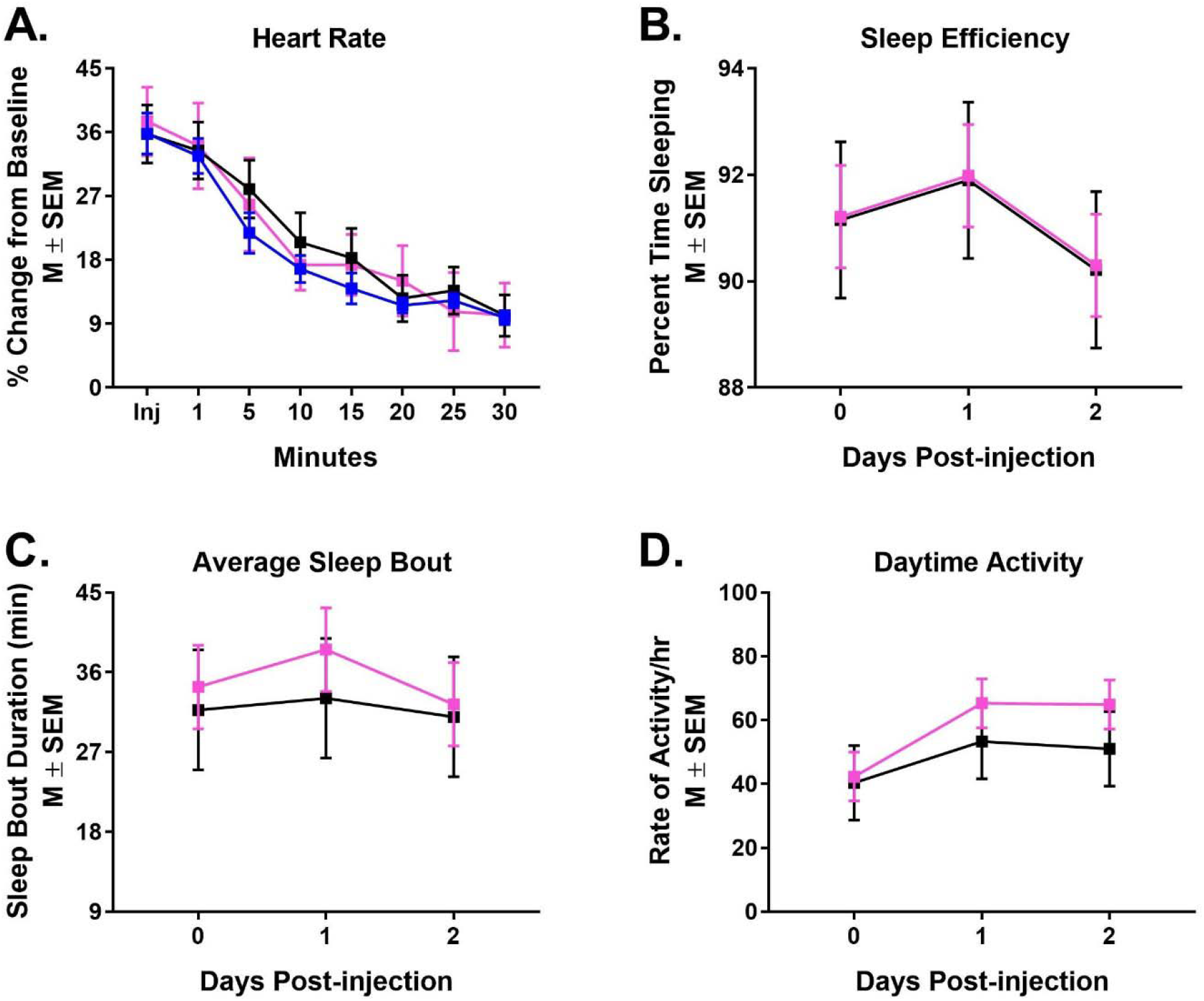
uPSEMs do not influence heart rate, sleep or activity levels. Percent change from baseline heart rate over 30 min of monitoring (A) after a s.c. injection of uPSEM817 (0.064 mg/kg; pink), uPSEM792 (0.87 mg/kg; blue) or vehicle (black). Sleep efficiency (B), average sleep bout duration (C), and rate of activity/hr during the day (D) after s.c. administration of uPSEM817 (0.064 mg/kg) or vehicle.

### Analysis of sleep and daytime activity

We investigated whether a 0.064 mg/kg dose of uPSEM817 might have an impact on sleep or daytime activity levels similar to that of varenicline.(Ashare et al., 2017; Savage et al., 2015) The sleep efficiency, latency of sleep onset, number of uninterrupted sleep bouts, and average duration of sleep bouts did not differ across days of observation (Day: F [2, 23] = 0.84, p = 0.44; F [2, 24] = 1.92, p = 0.17; F [2, 24] = 0.19, p = 0.83; F [2, 23] = 0.31, p = 0.74, respectively). uPSEM817 did not impact any of these sleep measures either on the day of injection, nor 1 or 2-days post-injection (Drug: sleep efficiency F [2, 23.9] = 0.0005, p = 0.98; latency of sleep onset F [1, 24] = 0.06, p = 0.80; number of uninterrupted sleep bouts F [1, 24] = 0.44, p = 0.51; average duration of sleep bouts F [1, 23.8] = 0.50, p = 0.49; Figure 2B & C). The rate of activity per hour during the day also remained the same (F [1, 24] = 1.31, p = 0.26; Figure 2D).

## DISCUSSION

Studies in mice have demonstrated that the uPSEMs activate their cognate PSAMs with low-nanomolar to sub-nanomlar potency and high selectivity (Magnus et al., 2019). The intracerebral bioavailability of systemically administered compounds can differ significantly between rodent and primate species (Syvänen et al., 2009), thus our demonstration that the uPSEMs have good brain penetrance, relatively slow washout (half-lifes of approximately 70 and 90 min for uPSEM817 and uPSEM792, respectively), and negligible conversion to varenicline or hydroxyvarenicline in rhesus monkeys suggests that they will be useful compounds for application to neuroscience studies in this species.

Two routes of administration were tested in our studies. We found similar results with i.v. and s.c. delivery, which would increase experimental flexibility when using these compounds for *in vivo* experiments in non-human primates (NHPs). The main difference was that i.v. delivery resulted in higher concentrations of the compounds. Both routes of administration resulted in stable concentrations (as measured in plasma and CSF) for 2-3 hours after administration. For both compounds, the concentrations achieved in brain are sufficient to activate PSAMs.

Both uPSEMs tested in the present study (uPSEM792 and uPSEM817) have structural similarity to varenicline, an FDA-approved smoking cessation drug. Our analyses did not detect metabolism to varenicline when using doses of 0.064 mg/kg uPSEM 817 or 0.87 mg/kg uPSEM792. Hydroxyvarenicline, an intermediate compound, was detected at low levels in the plasma after uPSEM817 administration, but there were negligibile concentrations in the CSF. The conversion of uPSEM817 to hydroxyvarenicline corresponds to de-alkylation of a propyl group. Hydroxyvareniciline was not detected in plasma or CSF after uPSEM792 administration, although due to technical difficulties detection of hydroxyvarenicline for the samples collected after PSEM792 assay (0.5 ng/mL) was less sensitive than the assay for the metabolite in uPSEM817 (0.1 ng/mL).

The doses used in the present study (0.064 mg/kg for uPSEM817, and 0.87 mg/kg for uPSEM792) were selected on the basis of the lowest effective doses measured in a mouse behavioral assay.(Magnus et al., 2019) The uPSEM792 dose was thus injected at a 14-fold higher dose than uPSEM817. Despite this difference, the recorded peak plasma concentration of uPSEM792 was ~50-fold higher than that of uPSEM817. This may reflect differences in transport into the brain and other tissues or a potential saturation of clearance mechanisms when we administered the higher uPSEM792 dose. In addition, uPSEM817 (cLogP: 2.9) is more hydrophobic than uPSEM792 (cLogP: 0.26), which may be associated with greater distribution to compartments outside of the plasma.

Effective brain access after systemic injection by crossing the blood brain barrier (BBB) is a requirement for *in vivo* neuroscience applications of chemogenetics. It has been shown previously that neither uPSEM is a substrate for the P-glycoprotein efflux pump (Magnus et al., 2019). Our results measuring CSF and parenchyma levels of uPSEM817 and uPSEM792 showed that these compounds have good brain bioavailability in monkeys. This is in agreement with previous rodent studies demonstrating brain penetrance of these compounds using *in vivo* positron emission tomography, in which binding of the radioligand ^18^F-ASEM to PSAM receptors was abolished by systemic injection of of uPSEM817 or uPSEM792 (Magnus et al., 2019).

Both uPSEMs had adequate BBB permeability as shown by their CSF:plasma ratios (uPSEM817 = 0.45 and uPSEM792 = 0.62). A previous study analyzing the pharmacokinetics of the structurally similar compound varenicline in rats reported a CSF:plasma ratio of 1.25 (Rollema, 2010), but this was based on the unbound fraction of the compound, which was not measured in our study. Therefore, the CSF:plasma ratios measured in our study, may be an underestimate.

The concentration of uPSEM817 (approx. 19 to 50 nM) and uPSEM792 (approx.150 to 250 nM) in the brain parenchyma recorded in our microdialysis experiments are more than sufficient to activate PSAMs, based on *in vitro* experiments in which the EC50s at PSAM^4^-GlyR channels were calculated as 0.3 and 2.3 nM for uPSEM817 and uPSEM792 respectively (Magnus et al., 2019). Notably, the parenchyma:CSF ratios were markedly different between these two compounds. The peak uPSEM817 concentration in the microdialysate (49.95nM) was approximately 8-fold higher than the peak concentration in the CSF (6.37nM). Conversely, the peak uPSEM792 concentration in the microdialysate (254.76nM) was approximately 65% of the peak concentration in CSF (405.76nM). This indicates that uPSEM817 may preferentially distribute to the parenchyma over the CSF. Several studies indicate that the parenchymal and CSF compartments can have distinct drug distribution properties (Aird, 1984; Liu, 2009). One caveat for the interpretation of the microdialysis results is the potential variability introduced by differences in the relative recovery correction factors for each microdialysis experiment; values reported for microdialysis are corrected values, based on a one-time recovery for the probe, done at the end of the experiment (when the probe may have reduced performance, possibly underestimating recovery). Also, microdialysis in the parenchyma measured the unbound fraction of drug, whereas CSF measurements were for total drug concentration. Although the concentrations measured in the brain are unlikely to activate nAChR receptors (Magnus et al., 2019), our results suggest that 5-fold to 10-fold lower doses of uPSEMs could be used for *in vivo* chemogenetic experiments in NHPs to minimize the possibility of off-target effects.

Plasma levels of the uPSEMs reached a maximal concentration 28 nM and 2 μM for intraveneously administered uPSEM817 and uPSEM792, respectively. At similar concentrations, *in vitro* testing has shown that uPSEM817 does not activate either α4β2- or α7-nAChRs. In contrast, uPSEM792 could activate endogenous α4β2-nAChR (EC50 0.52 μM), but endogenous α7-nAChRs are not activated when exposed to 2 μM of uPSEM792. However, neither uPSEM had significant effects on heart rate or sleep when compared to vehicle injections (fig, 2). Off-target side effects could influence behavioral neuroscience study results, as has been shown recently with other chemogenetic ligands (Ilg et al., 2018; MacLaren et al., 2016; Manvich et al., 2018; Upright and Baxter, 2020). While the search for off-target effects in our study was limited, the lack of identified adverse effects in this study and the evidence that considerably lower doses of uPSEMs are suitable for use in NHP, further increase the attractiveness of uPSEMs for neuroscience chemogenetic studies.

Although we did not examine the behavioral effects of modulating neuron activity in vivo with uPSEMs in NHPs, our previous work demonstrated that the latest generation of PSAMs are effective for modulating neuronal activity in the rhesus monkey brain *in* vivo (Magnus et al., 2019). The combination of good brain penetrance, slow washout, negligible conversion to psychoactive compounds, and absence of peripheral sideeffects at the doses tested in the present study leads us to conclude that uPSEM817 and uPSEM792 are well-suited for use in combination with PSAMs for applications in systems neuroscience.

## Acknowledgements

The authors would like to thank David Ji (B.S.) for his contribution to the Actical data analysis, Damien Pitard for his assistance with the awake heart rate monitoring experiments, Charalotta Campbell of Q2 solutions for LC-MS analysis, and Xing Hu for help in the microdialysis experiments.

## Funding

This work was funded by NIH-NINDS 1R21NS106346 (AG) and the NIH, Office of Research Infrastructure Program, P51 OD011132. MAGE, BJR, GPF and JYS were supported by the Intramural Research Program, NIMH, NIH, DHHS annual report ZIAMH 002619. Any opinions expressed are the authors’ own, and do not necessarily represent views of the NIMH, NIH, DHHS.

## Declarations of interest

Author S.M.S. has issued patents on this technology and owns stock in Redpin Therapeutics, LLC, which is a biotech company focusing on therapeutic applications of chemogenetics. S.M.S. is a cofounder and consultant for Redpin Therapeutics. All of the other authors (AG, JR, MAGE, BJR, GPF, JYS and TW) declare no competing financial interests.

## Supplemental Material

### Liquid chromatograph mass spectrometry (LC-MS)

Bioanalysis data for all compounds were obtained using an AB Sciex API 4000 triple quadrupole LC-MS device in positive ionization mode, coupled with a LEAP PAL HTS-xt autosampler and Shimadzu 30AD pumps. To achieve a lower limit of quantification for uPSEM817 plasma samples, data was obtained on an AB Sciex API 5500 triple quadrupole LC-MS under UPLC (ultra-performance liquid chromatography) conditions. Gradient information for each compound is listed in Supplementary Table 1.

uPSEM817 was monitored at a mass to charge ratio (m/z) of 270.30/199.20. For CSF samples, mobile phases consisted of 5mM ammonium bicarbonate in ultra-pure water (A) and 5 mM ammonium bicarbonate in methanol (B). A Sprite ECHELON C18 4μm 20 x 2.1mm column used for chromatography. Sample injection volume was set at 10uL with an approximate run time of 0.72 min. For plasma samples, mobile phases of 0.1% formic acid in ultra-pure water (A) and 0.1% formic acid in acetonitrile (B). An Acquity UPLC BEH C18 1.7 μm, 2.1 × 50 mm column was used for chromatography. Sample injection volume was set at 15 μL with an approximate run time of 1.5 min. Lower limit of quantification (LLOQ) for uSPEM817 was 0.1 ng/ml for plasma and CSF samples, and 0.25 ng/ml for microdialysate samples.

Hydroxyvarenicline was monitored at an a m/z of 228.10/199.10 and varenicline was monitored at an m/z of 212.10/183.20. Mobile phases consisted of 0.4% trifluoroacetic acid (TFA) and 1 mM ammonium bicarbonate in ultra-pure water (A) and 0.4% TFA and 1 mM ammonium bicarbonate in acetonitrile (B). A Sprite ECHELON C18 4μm 20×2.1mm column was used for chromatography. Sample injection volume was set at 10 μL with an approximate run time of 0.72 min. LLOQ for hydroxyvarenicline was between 0.1 to 0.5 ng/ml.The lowest limit of detection for varenicline was 0.1 ng/ml.

uPSEM792 was monitored at an m/z of 242.20/213.20. Mobile phases consisted of 0.4% TFA and 1 mM ammonium bicarbonate in ultra-pure water (A) and 0.4% TFA and 1 mM ammonium bicarbonate in acetonitrile (B). A Sprite ECHELON C18 4μm 20 x 2.1mm column was used for chromatography. Sample injection volume was set at 10 μL with an approximate run time of 0.72 min. LLOQ for uPSEM792 was 0.25 ng/ml for plasma and 0.1 ng/ml for CSF samples.

Standard curves for each test compound ranged from 0.1 to 5000 ng/mL, with quadratic 1/x2 weighting, and all data were acquired using Analyst v1.6.3. A summary of mass spectrometer parameters, including Internal Standards (IS) along with LC conditions can be found in supplementary table 2.

### Heart Rate Monitoring

For monitoring via the wearable heart rate sensors (FitBit Inspire 2, San Francisco, CA or Polar OH1, Polar Electro Oy, Kempele, Finland), the hair was shaved from around the monkeys leg and the optical sensor was placed in the bend of the knee and secured in place using elastic disposable bandaging (Coban Self-Adherent Wrap, 3M Healthcare, St Paul, MN). The accuracy of the wearable optical heart rate sensors was validated in anesthetized and awake monkeys. During the uPSEM817 s.c. administration challenge, sedated monkeys were monitored via an optical pulseoximeter (SurgiVet® Advisor® Vital Signs Monitor Dublin, OH) placed on the animal’s ear and a wearable optical heart rate monitor (Fitbit Inspire 2 or Polar OH1) in the bend of their knee. To ensure that readings were also accurate under awake conditions, one animal was monitored with both devices during a vehicle injection. Regardless of sedation or awake conditions, the wearable optical heart rate sensors proved to be 99.66% accurate compared to the optical pulse oximeter (SurgiVet® Advisor® Vital Signs Monitor; Supplementary Table 3).

**Supplementary Table 1.**
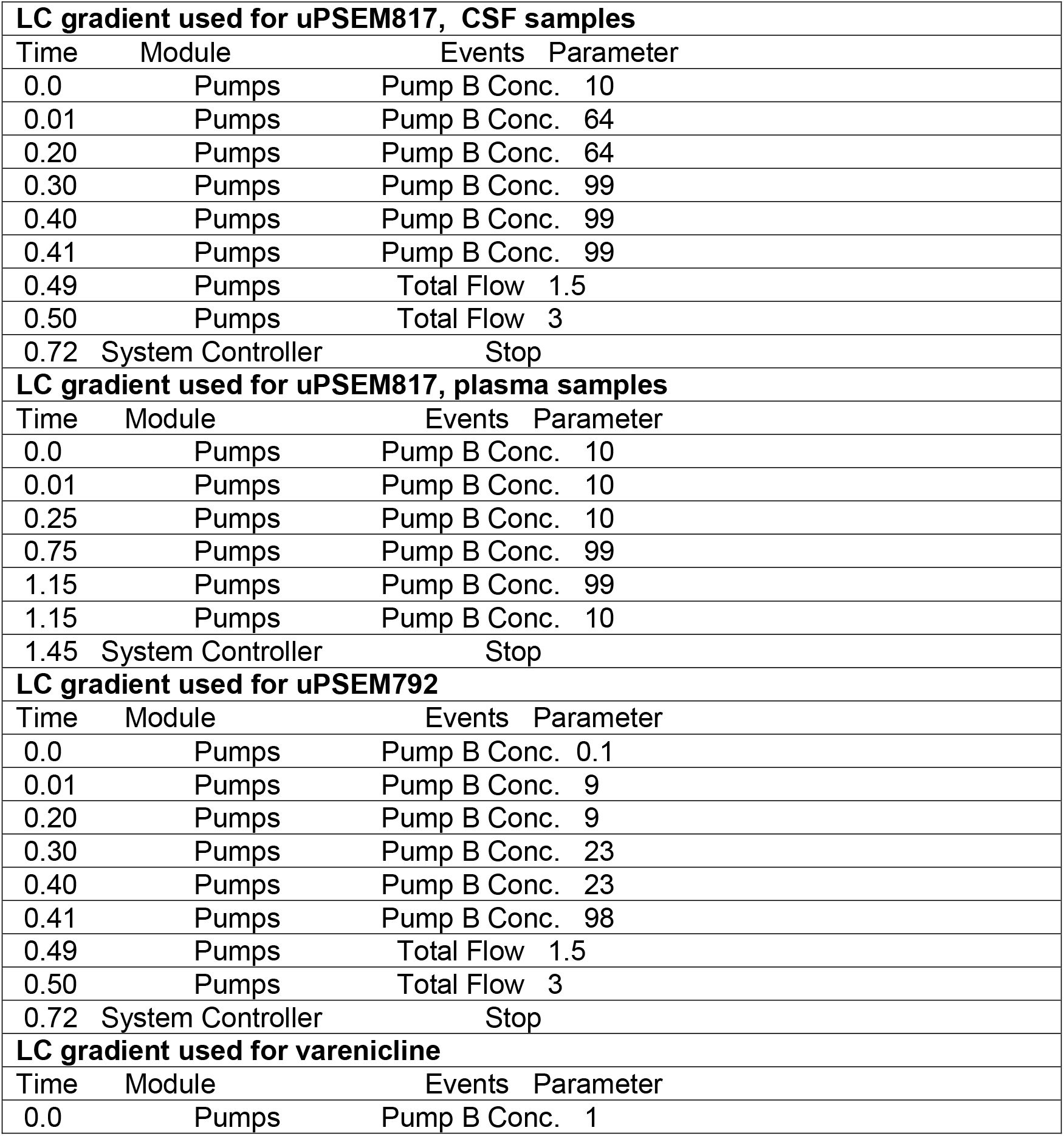
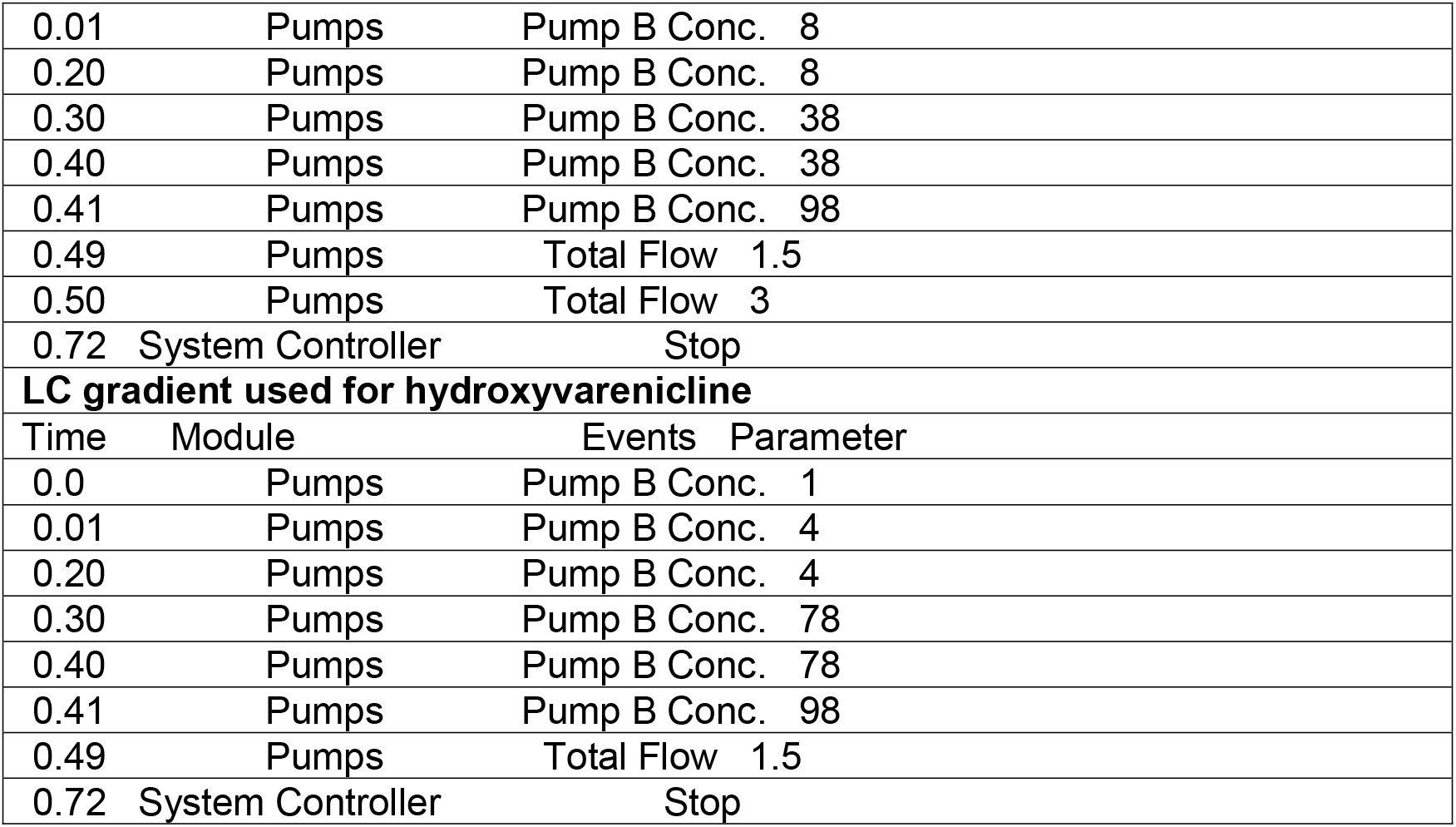
LC gradients used for analysis of uPSEM and metabolites

**Supplementary Table 2.**
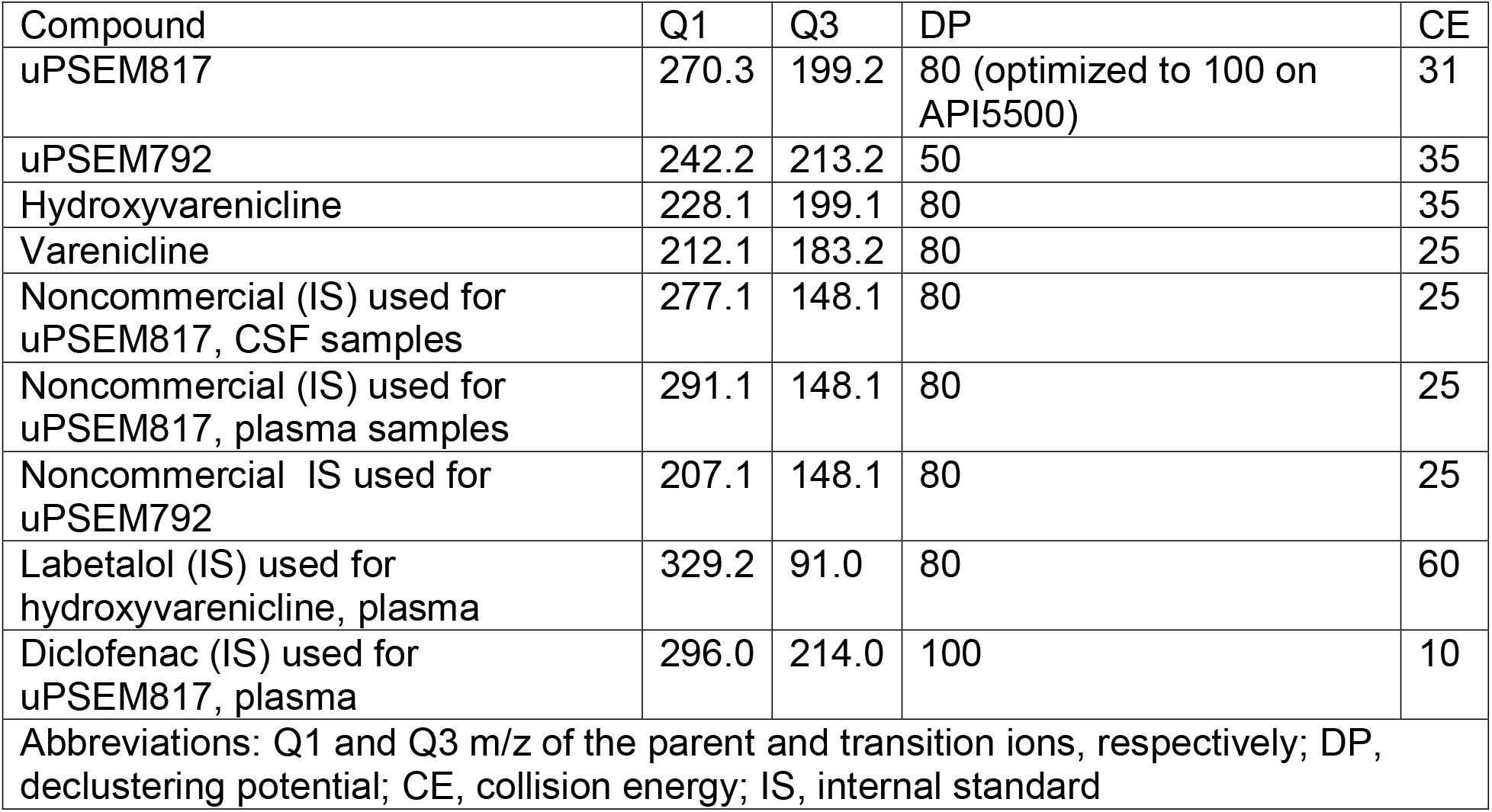
Mass Spectrometer Parameters used for analysis

**Supplementary Table 3.**
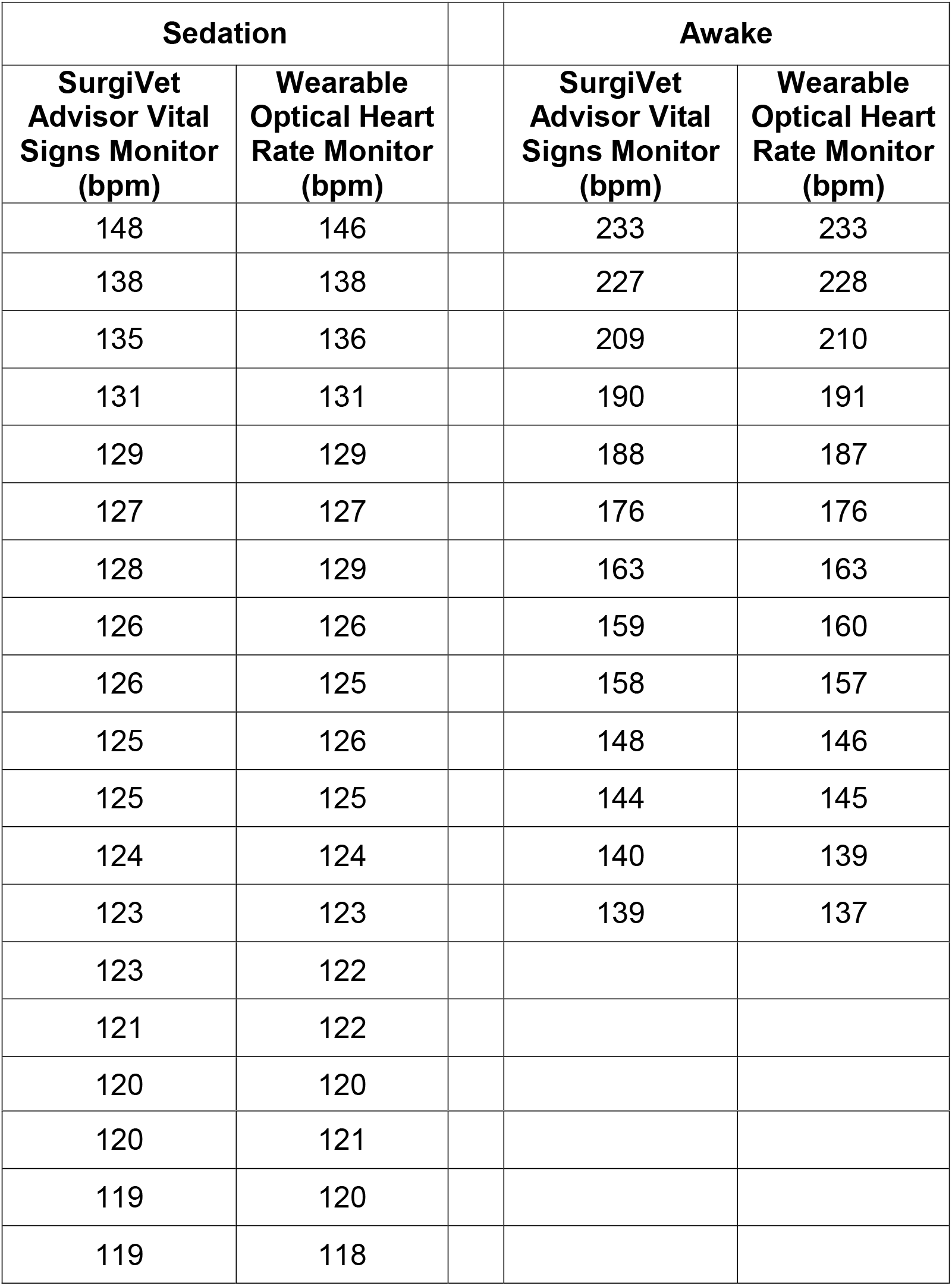
Correspondence between optical pulse-oximeter devices

| Sedation |  |  | Awake |  |
| --- | --- | --- | --- | --- |
| **SurgiVet**<br>**Advisor Vital**<br>**Signs Monitor**<br>**(bpm)** | **Wearable**<br>**Optical Heart**<br>**Rate Monitor**<br>**(bpm)** |  | **SurgiVet**<br>**Advisor Vital**<br>**Signs Monitor**<br>**(bpm)** | **Wearable**<br>**Optical Heart**<br>**Rate Monitor**<br>**(bpm)** |
| 148 | 146 |  | 233 | 233 |
| 138 | 138 |  | 227 | 228 |
| 135 | 136 |  | 209 | 210 |
| 131 | 131 |  | 190 | 191 |
| 129 | 129 |  | 188 | 187 |
| 127 | 127 |  | 176 | 176 |
| 128 | 129 |  | 163 | 163 |
| 126 | 126 |  | 159 | 160 |
| 126 | 125 |  | 158 | 157 |
| 125 | 126 |  | 148 | 146 |
| 125 | 125 |  | 144 | 145 |
| 124 | 124 |  | 140 | 139 |
| 123 | 123 |  | 139 | 137 |
| 123 | 122 |  |  |  |
| 121 | 122 |  |  |  |
| 120 | 120 |  |  |  |

|  |  |
| --- | --- |
| 120 | 121 |
| 119 | 120 |
| 119 | 118 |

